# AngleCam V2: Predicting leaf inclination angles across taxa from daytime and nighttime photos

**DOI:** 10.1101/2025.09.17.676742

**Authors:** Luis Kremer, Jan Pisek, Ronny Richter, Julian Frey, Daniel Lusk, Christiane Werner, Christian Wirth, Teja Kattenborn

## Abstract

1. Understanding how plants capture light and maintain their energy balance is crucial for predicting how ecosystems respond to environmental changes. By monitoring leaf inclination angle distributions (LIADs), we can gain insights into plant behaviour that directly influences ecosystem functioning. LIADs affect radiative transfer processes and reflectance signals, which are essential components of satellite-based vegetation monitoring. Despite their importance, scalable methods for continuously observing these dynamics across different plant species throughout day-night cycles are limited.
2. We present AngleCam V2, a deep learning model that estimates LIADs from both RGB and near-infrared (NIR) night-vision imagery. We compiled a dataset of over 4,500 images across 200 globally distributed species to facilitate generalization across taxa. Moreover, we developed a method to simulate pseudo-NIR imagery from RGB imagery to enable an efficient training of a deep learning model for tracking LIADs across day and night. The model is based on a vision transformer architecture with mixed-modality training using the RGB and the synthetic NIR images.
3. AngleCam V2 achieved substantial improvements in generalization compared to AngleCam V1 (R^2^ = 0.62 vs 0.12 on the same holdout dataset). Phylogenetic analysis across 100 genera revealed no systematic taxonomic bias in prediction errors. Testing against leaf angle dynamics obtained from multitemporal terrestrial laser scanning demonstrated the reliable tracking of diurnal leaf movements (R^2^ = 0.61-0.75) and the successful detection of water limitation-induced changes over a 14-day monitoring period.
4. This method enables continuous monitoring of leaf angle dynamics using conventional cameras, enabling applications in ecosystem monitoring networks, plant stress detection, interpreting satellite vegetation signals, and citizen science platforms for global-scale understanding of plant structural responses.

## 1 Introduction

Some plants hold their leaves horizontally like umbrellas, while others keep them upright like rows of solar panels. This difference in orientation, ranging from flat to erect, is described as the vertical leaf angle. Plants have evolved a wide variability of vertical leaf angle orientations across species, growth forms, and geographic regions (Puglielli et al., 2024).

Even within a single plant, leaf angles are not uniform: upper canopy leaves often stand more erect, channeling light deeper into the crown, while lower leaves spread out more horizontally to intercept the remaining light (Niinemets, 2010). This variation has long been recognized, with Charles Darwin already emphasizing the ecological and physiological relevance of leaf angles in *The Power of Movement in Plants* (Darwin & Darwin, 1883). Since then, leaf angles have been shown to regulate canopy light capture, drive competition, and even influence microclimates and productivity (Hikosaka & Hirose, 1997; Mullen et al., 2006; Niinemets, 2010).

The vertical orientation of individual leaves is commonly described by their surface angle, ranging from 0° (horizontal) to 90° (vertical). The variability of these angles within a canopy is referred to as the leaf inclination angle distribution (LIAD). Importantly, LIADs are not static: Plants dynamically adjust leaf orientations in response to diurnal rhythms, phenological changes, and environmental stimuli such as light availability, temperature, competition, and water stress (Geldhof et al., 2021). These adjustments include nastic movements (non-directional responses to stimuli like light and temperature), tropisms (directional growth responses), and passive or active responses, such as leaf drooping or wilting due to drought or heat (Apelt et al., 2017; Niinemets, 2010; Puglielli et al., 2017). Moreover, shifts in leaf angle can serve as rapid indicators of stress conditions such as heat or drought, or as protective responses to excessive radiation and photoinhibition (Ehleringer et al., 1987; Kattenborn et al., 2022; Van Zanten et al., 2010; Werner et al., 2001), thereby altering canopy energy balances (Leuzinger & Körner, 2007; Sastry et al., 2018). Leaf angle variations also affect how light is reflected to satellite sensors, which is crucial for global vegetation monitoring (Braghiere et al., 2021; Dechant et al., 2020; Jablonski et al., 2025; Kattenborn et al., 2024).

Despite their importance, few scalable techniques exist for automatically tracking dynamic changes in LIADs (Yang et al., 2023). While inertial measurement units (IMUs) can track individual leaf movements (Geldhof et al., 2021), they are not scalable to entire canopies. 3D point cloud-based approaches using photogrammetry (Qi et al., 2019), terrestrial laser scanning (Bailey & Mahaffee, 2017; Murithi et al., 2025; Stovall et al., 2021; Zheng & Moskal, 2012), or stereo vision (Bernotas et al., 2019; Biskup et al., 2007; Müller-Linow et al., 2015) show promise but require specialized hardware and calibration, limiting their usability in field conditions.

Yet humans can easily perceive differences in leaf angles from real-life observations or photographs (Pisek et al., 2011; Zou et al., 2014). Automating this task using computer vision techniques led to the development of AngleCam (Kattenborn et al., 2022, 2024), a deep learning model designed to estimate LIADs from single, horizontally oriented RGB photographs or video frames. AngleCam was trained on approximately 2500 images and corresponding leaf inclination angle distributions, which were derived from annotating individual leaves in the image frames, and compared against independent TLS-derived LIADs across 25 plant species. The plausibility and potential of AngleCam were underlined, given that predicted leaf inclination angle distributions over multiple months had a tight relationship to environmental conditions, such as radiation, temperature, or soil humidity. Despite these potentials, the original AngleCam (V1) remains limited to daylight observations and has restricted generalizability due to the limited diversity of its training dataset.

Here, we present AngleCam V2, which addresses these limitations by expanding the training and validation dataset to over 4500 images across 200 species with broad geographic coverage. The increased taxonomic and environmental diversity enhances model generalization across diverse plant communities and scene conditions. A fundamental advancement is the capability to process both RGB and near-infrared (NIR) night-vision imagery, enabling continuous monitoring of leaf angle dynamics across complete diurnal cycles. We validate the model on a holdout dataset and assess its generalization across genera. Additionally, we test the approach in controlled indoor experiments, which include a comparison with multitemporal terrestrial laser scanning (TLS) and monitoring of a plant under water limitation. The enhanced generalization and temporal capabilities could make this approach a promising candidate for ecosystem monitoring networks and citizen science applications, with potential for contributing to global-scale understanding of leaf angle patterns and their environmental drivers.

## 2 Materials and Methods

### 2.1 AngleCam model development

#### 2.1.1 RGB image data across taxa and geographic regions

The dataset for model training and validation grew from 2680 to 4801 images, enhancing taxonomic and morphological diversity with AngleCam V2 (Fig. 1). Unlike AngleCam V1, which relied solely on two datasets from 2021, AngleCam V2 includes these plus three additional collections (Tab. 1). The first is a diverse set from the Leipzig area, featuring 100 species from various environments. The second set consists of image time series taken within the project at the Canopy Crane research platform (Leipzig Canopy Crane [LCC]) of the German Centre for Integrative Biodiversity Research (iDiv) Halle-Jena-Leipzig. This site covers a Leipzig floodplain forest, a structurally complex hardwood forest dominated by tree species such as European ash (*Fraxinus excelsior* L.), English oak (*Quercus robur* L.), Sycamore maple (*Acer pseudoplatanus* L.), European hornbeam (*Carpinus betulus* L.), and small-leaved lime (*Tilia cordata* Mill) (see Richter et al., 2022, for details). Together, both datasets provided a total of 102 different species. While *Tilia cordata* and *Acer pseudoplatanus* represented approximately 50% of the dataset, this reflects extensive temporal sampling from continuous monitoring at the Leipzig Canopy Crane, capturing a large variability of LIADs of these species across diverse environmental conditions and seasonal phenology rather than simple taxonomic overrepresentation (Fig. 1). These image series were captured using TLC-200 Pro timelapse cameras (Brinno Inc., Taipei, Taiwan) in high-dynamic range (HDR) mode (further details see Kattenborn et al., 2022).

**Figure 1:**
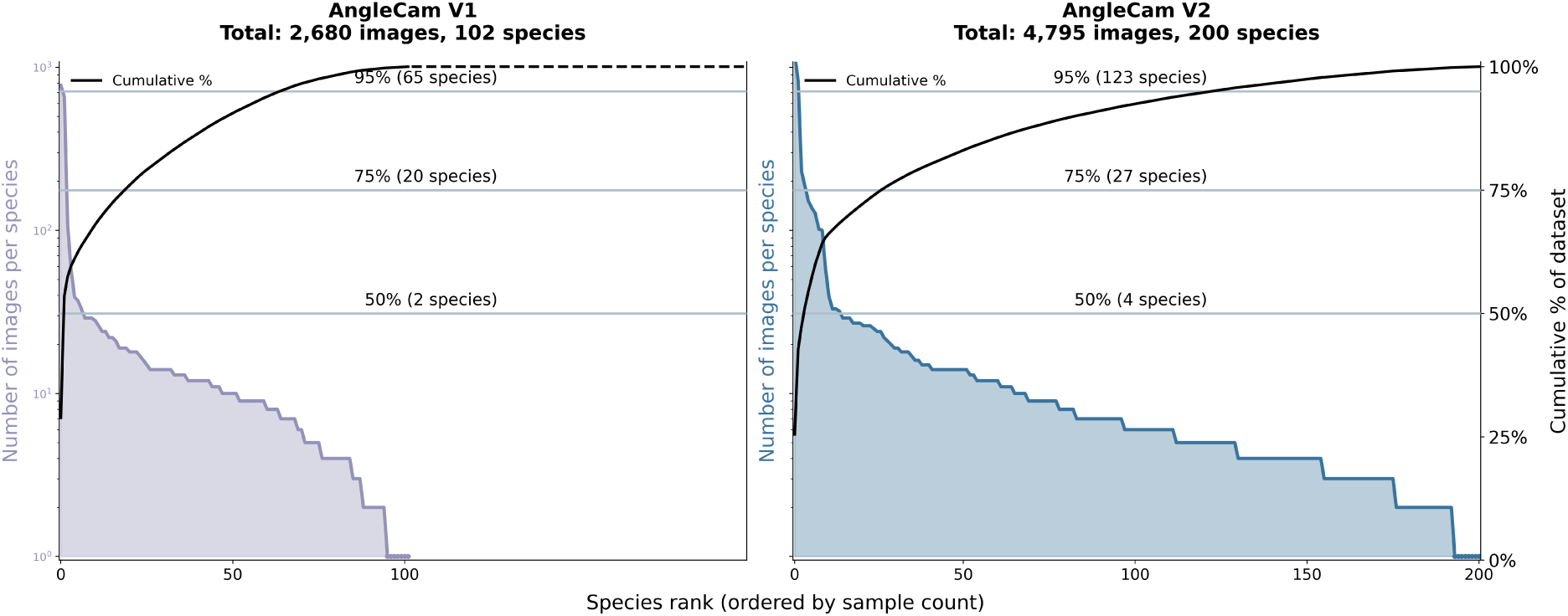
Species distribution in AngleCam (V1 & V2) training datasets. Rank-abundance curves showing the number of images per species (left y-axis, log scale) and cumulative dataset coverage (right y-axis) for Version 1 (left panel, 2680 images across 102 species) and Version 2 (right panel, 4801 images across 200 species). The curves illustrate how images are distributed across species in each dataset, with reference lines indicating the number of species required to achieve 50%, 75%, and 95% of total dataset coverage.

**Table 1:** Datasets used for AngleCam V1 and V2 model training and validation.

| Dataset | Year(s) | Species Count | Image Count | Description | AngleCam Version |
| --- | --- | --- | --- | --- | --- |
| Area of Leipzig | 2021 | 100 | 1237 | Natural/semi-natural areas, parks, gardens, indoor plants | V1 & V2 |
| Leipzig Canopy Crane (LCC) | 2021 | 2 | 1431 | Floodplain forest, a hardwood ecosystem | V1 & V2 |
| MyDiv experiment | 2022 | 9 | 1422 | Tree monocultures | V2 only |
| Global Collection | 2012–2022 | >100 | 676 | Botanical gardens, arboreta worldwide | V2 only |
| Indoor experiments | 2024–2025 | 3 | 35 | Controlled indoor plant monitoring | V2 only |

With the new version of AngleCam, we have expanded the training and validation dataset to make the model more robust against various leaf shapes and illumination conditions. The entire dataset now includes three additional datasets in addition to the two from the first version:

- Image series taken within the field experiment MyDiv (Mycorrhiza in tree Diversity effects on ecosystem functioning; Ferlian et al., 2018)
- A globally sourced collection of over 600 tree and shrub samples representing more than 100 species from diverse climates and biomes.
- An independent test dataset with image series of three indoor plant species, including two validated against terrestrial laser scans (described in Section 2.3 & 2.4)

The MyDiv experiment, conducted at the Bad Lauchstädt Experimental Research Station of the Helmholtz Center for Environmental Research-UFZ in Saxony-Anhalt, Germany, is structured to investigate the effects of tree diversity on ecosystem functions (Ferlian et al., 2018). The data for this analysis specifically used measurements from monoculture plots of nine temperate deciduous tree species: *Acer pseudoplatanus* L., *Aesculus hippocastanum* L., *Fraxinus excelsior* L., *Sorbus aucuparia* L., *Betula pendula* R., *Carpinus betulus* L., *Fagus sylvatica* L., *Quercus petraea* (Matt.) Liebl., and *Tilia platyphyllos* Scop. For each of these nine species, two time-lapse cameras were deployed in the plot center of the respective monoculture plots. The cameras were positioned to capture a horizontal field of view of branches from two individuals located centrally within the plot. The installation height corresponded to the upper third of the tree crowns. Time series images were collected at 5-minute intervals during the primary vegetation period from June 6 to October 5, 2022 (further details see Kattenborn et al., 2024).

To improve the model’s robustness across genera and scene conditions, we integrated a comprehensive dataset that includes 676 horizontal plant photographs representing over 100 distinct species across a broad taxonomic and ecological spectrum. The observations were collected from botanical gardens and arboreta worldwide, including locations in Europe (Czech Republic, Estonia, France, Italy, Netherlands, Portugal, Spain, Sweden, and the United Kingdom), as well as in the USA, Israel, and Australia. This collection covers a variety of biomes and climates, ranging from Mediterranean and temperate to subtropical regions. For several species and genera, such as *Eucalyptus*, *Quercus*, and *Betula*, multiple samples from diverse locations were collected, resulting in a dataset that includes both intra- and interspecific variation under varied environmental conditions.

#### 2.1.2 Labeling RGB images with leaf inclination angle distributions

Deep learning models benefit from large, well-annotated training datasets. To generate leaf angle references efficiently, we used visual estimation based on RGB images. Previous studies obtained estimates of leaf inclination angle distributions by measuring the vertical angle of leaves that are oriented perpendicular to the camera’s line of sight (Pisek et al., 2011; Ryu et al., 2010). Multiple measurements of individual leaves can be summarized as a leaf inclination angle distribution. However, these strict geometric requirements limit the number of usable samples per image, the robustness of a leaf inclination angle distribution, and pose challenges for species with complex or curled leaf forms (Murithi et al., 2025). Here, we overcome these limitations by estimating the average leaf inclination of whole leaf surfaces through visual interpretation of their apparent angle from horizontal in RGB images. This approach was already successfully applied for AngleCam V1 (Kattenborn et al., 2022).

We obtained a LIAD for each reference image by sampling the average leaf surface inclinations of 20 leaves, which provided robust estimates while allowing efficient annotation (Kattenborn et al., 2022). An evenly spaced point grid was used to ensure balanced sampling. The 20 leaf samples were converted into probability distributions to account for estimation inaccuracies across the full 0°–90° leaf angle range. We employed the two-parameter beta distribution, which is well-suited to model various leaf inclination angle distribution shapes from strongly horizontal (planophile) to uniform, spherical, or strongly vertical (erectophile) forms (Goel & Strebel, 1984). Beta distribution fitting was performed in Python using the scipy.stats module (v1.13.0), with parameters (*α*, *β*) estimated by maximum likelihood.

We trained the model to predict the entire probability density function (PDF) across the 0°–90° range at 2° intervals, rather than directly predicting the (*α*, *β*) parameters. This allows the model to deviate from an idealized beta distribution. To enhance generalization, we perturbed the fitted (*α*, *β*) values of the reference LIADs, generating 50 synthetic variants for each image by sampling (*α*, *β*) within *±*20% of the standard deviation of the maximum likelihood estimates. During training, one randomly selected augmented distribution was used as the target for each image per epoch, while the original unperturbed distribution served as the reference for validation. This produced reference data for 4801 RGB images.

#### 2.1.3 Generating training data on pseudo-NIR images

While AngleCam V1 was trained exclusively on RGB images, we specifically adapted AngleCam V2 to also process NIR images acquired under nighttime conditions, enabling fully continuous monitoring of leaf angle dynamics.

Night-vision images are captured by cameras with infrared light-emitting diodes (NIR-LEDs) that emit near-infrared radiation (850-940 nm). The camera sensor records reflected NIR light, resulting in grayscale images that represent backscatter intensity (Sun et al., 2023). These images differ from daylight RGB images in several ways: the spectrum varies (e.g., vegetation appears bright in NIR), the effective range of the NIR LED is around 9 meters, and illumination decreases with the square of the distance from the light source (Pharr et al., 2023).

These fundamental differences posed a challenge, as a model trained only on RGB images would perform poorly on NIR images due to modality mismatch. Given that a large dataset of labeled RGB data was already available (Tab. 1), we decided not to invest the same workload to create an equally sized dataset of labeled NIR imagery. Instead, we developed an approach to create synthetic NIR imagery from the already extensively labeled RGB data.

Firstly, we randomly converted 50% of the RGB in the training dataset to grayscale using luminance weighting (0.299R + 0.587G + 0.114B) to reduce reliance on color. Secondly, we randomly added a distance-based dimming to 50% of the grayscale images (pseudo-NIR images) to mimic the limited range of the NIR illuminator and the corresponding brightness falloff with distance (Fig. 2). For this, we scaled the pixel intensities of each grayscale image according to the pixel-wise distances estimated from computer vision–derived depth maps (Depth Anything V2; see details below). The final transformation was defined as:

**Figure 2:**
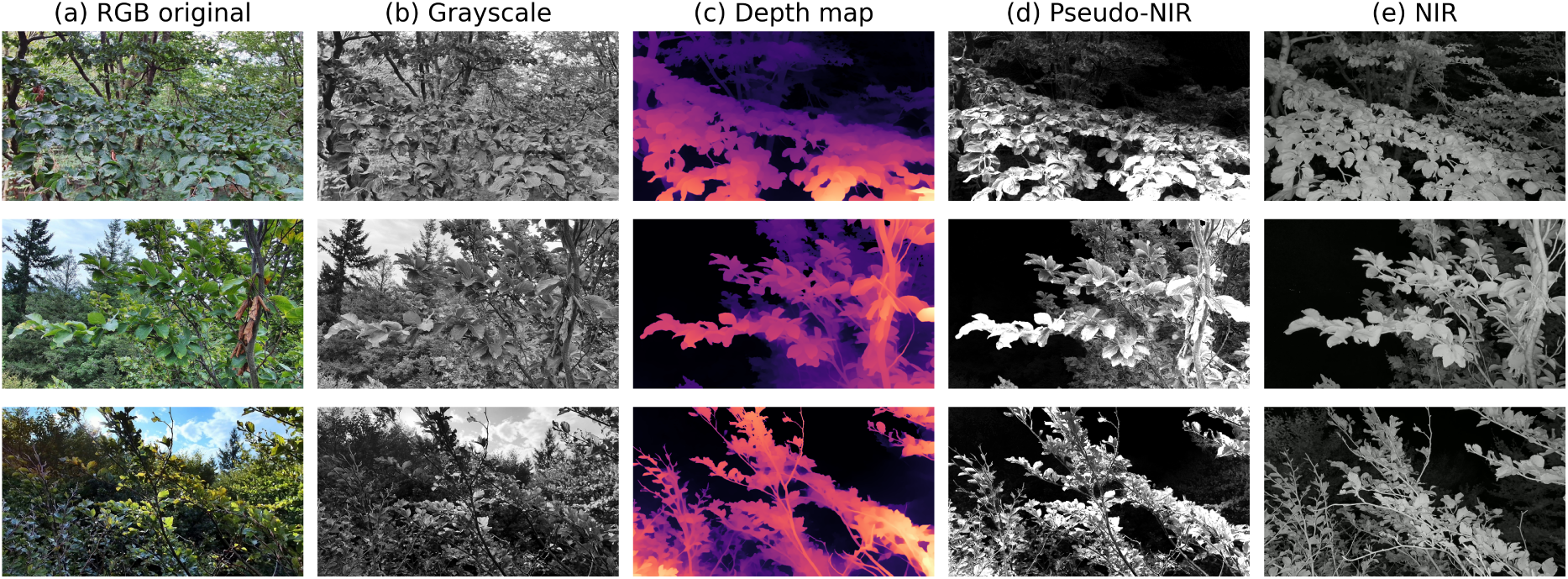
Pseudo-NIR image generation pipeline. The transformation from RGB to pseudo-NIR involves several steps: (a) RGB original image, (b) conversion to grayscale, (c) depth map estimation using Depth Anything V2 (closer objects to the camera appear brighter), and (d) pseudo-NIR image generation by scaling the intensity of the grayscale image with the corresponding depth map, simulating NIR-LED illumination falloff. (e) Actual NIR image captured at nighttime of the same scene.

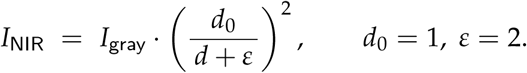

where *I*_gray_ is the grayscale image, *d* is the per-pixel depth (in meters), and *d*_0_ is a normalization distance. We set *ε* = 2 to ensure numerical stability for minimal depths and to slightly compress the distance attenuation. This yields pseudo-NIR images that appear brighter and less contrast-extreme (Fig. 2).

The depth maps used to simulate the NIR imagery were generated using Depth Anything V2, a state-of-the-art monocular depth estimation model (Yang et al., 2024). We used the large model variant, fine-tuned for metric depth estimation on the Virtual KITTI 2 dataset (Cabon et al., 2020). To align predicted depth values with the operational range of our NIR camera system, we constrained maximum depth predictions to 9 meters.

#### 2.1.4 Model architecture and training

For AngleCam V2, we revised the architecture by replacing the EfficientNet-B7 backbone with the self-supervised DINOv2 ViT-S/14 model as the feature extractor (Oquab et al., 2023). Vision transformers offer a comprehensive feature representation, effectively recognizing structures and capturing depth cues through their global context processing (Dosovitskiy et al., 2020; Tolan et al., 2024).

The DINOv2 ViT-S/14 backbone outputs a 384-dimensional embedding, which we passed into a lightweight regression head: a fully connected layer (384 → 128) with dropout (rate = 0.4), GELU activation (Hendrycks & Gimpel, 2016), another fully connected layer (128 → 43), and a softmax output layer for a 43-bin probability distribution across 2° intervals from 0° to 90°.

Images were resized to 224×224 pixels via linear interpolation. The augmentation pipeline included *RandomResizedCrop* (scale = [0.8, 1.0]) and horizontal flipping (50% probability), while photometric augmentations adjusted brightness and contrast (variation factors = 0.3) and added Gaussian noise (*σ* = 0.02) for improved robustness to sensor noise and artifacts.

Instead of training separate models for RGB and NIR imagery, we developed AngleCamV2 using a mixed-modality approach. This design ensures that, regardless of whether NIR or RGB data are available, no model switching or dataset separation is required. During each epoch, 50% of images were randomly presented as original RGB, while the remaining 50% were non-RGB variants: true grayscale and pseudo-NIR images generated through distance-based dimming (Section 2.1.3; Fig. 2). All images were normalized using ImageNet statistics (mean = [0.485, 0.456, 0.406], std = [0.229, 0.224, 0.225]).

The model was optimized using the AdamW optimizer with an initial learning rate of 1 *×* 10*^−^*^4^, weight decay of 0.01, and gradient clipping (max norm = 3.0, *L*_2_ norm type) to prevent overfitting (Loshchilov & Hutter, 2017). Learning rate scheduling was applied via *ReduceLROnPlateau* (reduction factor = 0.5, patience = 5 epochs, min learning rate = 1 *×* 10*^−^*^6^). Training lasted for 50 epochs with a batch size of 32, and Huber loss was used to reduce sensitivity to outliers (Sun et al., 2020). The final model was selected based on the lowest validation loss.

Prior to splitting the dataset, three RGB images per genus were extracted to assess phylogenetic prediction errors (see next Section 2.2). The remaining labeled data were split 80/20 into training and validation sets, resulting in approximately 3600 images for training and 900 for validation. The random split resulted in some overlap of species and acquisition sites between training and validation sets. While fully independent validation datasets would be ideal, creating stratified splits across multiple dimensions (species, sites, temporal series, and leaf angles) would result in impractically small subsets. Importantly, images within shared species were captured across different seasons, phenological stages, and environmental conditions, providing substantial variability. Validation was performed exclusively on original RGB images since the pseudo-NIR and grayscale variants share the same labels. The model’s capability to process NIR imagery was independently validated through controlled experiments using terrestrial laser scanning as reference data (Section 2.3).

### 2.2 Phylogenetic error analysis

Predictive models in ecology often exhibit systematic biases when applied across different taxonomic groups, particularly when training data is not balanced across taxonomic and phylogenetic lineages (Morales-Castilla et al., 2024). Such phylogenetic autocorrelation in prediction errors can indicate dataset gaps, model limitations, or evolutionary constraints that limit generalizability (Roberts et al., 2017). This consideration is relevant for our dataset, where temporal monitoring at specific sites resulted in uneven species representation, with approximately 50% of training images originating from four species. Given this taxonomic imbalance, we assessed whether the residuals of AngleCam V2 are phylogenetically autocorrelated.

For images where species identification was not available in the original metadata, we determined plant species using the Pl@ntNet online identification tool (PlantNet, 2025). To evaluate phylogenetic error in the residuals, three random images per genus were excluded from the training dataset and used for analysis. Since Pl@ntNet has slight identification uncertainties at the species level, the analysis was performed at the genus level. Genera with fewer than three samples were not analyzed due to insufficient significance. Consequently, from a total of 129 genera, 100 genera with 300 samples were included in the analysis.

For each genus, we calculated the LIAD residuals as the mean absolute deviation between AngleCam V2 predictions and labeled LIAD references. The genus-level errors were then mapped onto a phylogenetic tree to visualize error patterns between taxonomic groups. We quantified the phylogenetic signal in prediction errors using Pagel’s *λ*, which measures the degree to which closely related genera exhibit similar prediction errors (Pagel, 1999). A *λ* near 1 indicates a strong phylogenetic influence, while a *λ* near 0 suggests independence from phylogeny (Kamilar & Cooper, 2013). We estimated Pagel’s *λ* with the function *phylosig* of the *R*-package *phytools* (v.2.4-4).

We quantified phylogenetic autocorrelation in model residuals using Moran’s *I* statistic calculated on phylogenetic distances (Gittleman & Kot, 1990; Moran, 1950). Significant positive autocorrelation would indicate that prediction errors are clustered within particular clades, suggesting systematic biases that could limit model transferability to underrepresented taxonomic groups (Hawkins, 2012). We estimated Moran’s *I* with the function *moran.idx* of the *R*-package *adephylo* (v.1.1-17).

### 2.3 Multitemporal model evaluation of AngleCam V2 with TLS-derived leaf inclination angle distributions

#### 2.3.1 Experimental setup and data acquisition

We validated diurnal leaf inclination angle distributions obtained from AngleCam V2 against independent LIADs from terrestrial laser scanning (TLS). An indoor experiment was conducted with two species known for strong diurnal movements, *Maranta leuconeura* E.Morren and *Calathea ornata* (Linden) Körn, monitored from 17–20 January 2025.

The TLS scanner (Riegl VZ-400i; RIEGL Laser Measurement Systems GmbH, Horn, Austria) was positioned 1.5–2.0 m from the plants for unobstructed coverage, acquiring scans every 30 minutes. It operates with a near-infrared laser (1550 nm) and offers a measurement precision of 5 mm, with a field of view of 100° vertical and 360° horizontal.

Two Ubiquiti UniFi G5 Bullet cameras (Ubiquiti Inc., New York, NY, USA) were installed 1 m from each plant, capturing images every 30 minutes in both RGB and near-infrared modes (NIR range: 9 m) and synchronized with TLS acquisitions. Images were geometrically calibrated to correct for lens distortion.

#### 2.3.2 TLS data processing and AngleCam comparison

The TLS point clouds were processed to extract LIADs for comparison with AngleCam V2 predictions. The pipeline focused solely on leaf surfaces by manually defining bounding boxes to isolate target plants, eliminating stems, petioles, and background objects.

Point clouds were centered around the plant base for a consistent coordinate system. Surface normals were estimated using principal component analysis (PCA) within a 1 cm radius, identifying the direction of least variance (Jolliffe, 2002; Rusu et al., 2008). Normals were oriented toward the scanner for consistent angle calculations, and the point clouds were subsampled with a voxel grid at a 1 cm resolution to reduce density variation.

For statistical analysis, TLS-derived LIADs were temporally aligned with AngleCam predictions based on acquisition timestamps. Quantitative comparisons were performed on average leaf angle values from LIAD distributions and the closest-in-time AngleCam predictions, calculating metrics like the coefficient of determination (R^2^) and root mean square error (RMSE) for each matched time point.

### 2.4 Multitemporal model evaluation of AngleCam V2 under water limitation

We performed an additional indoor experiment to assess whether AngleCam V2 captures extended, multi-day trends under plant water limitation. We positioned an individual *Aglaonema commutatum* Schott in front of a window and recorded images as described above over 14 days (17–31 December 2024). Images were acquired every two minutes, resulting in 8783 images. The plant was last watered on December 12 and thus remained without irrigation for 19 days by the end of monitoring.

## 3 Results

### 3.1 Model performance on training and validation data

The model evaluation revealed distinct performance patterns across training and validation datasets, as well as a substantial improvement over the previous model version V1 (Figure 3). For the training dataset, we found a strong correspondence between the predicted and reference average leaf angles (R^2^ = 0.75, RMSE = 7.37°, n = 3591). The regression relationship indicated a minor overestimation of lower shallow angles and an underestimation of steeper angles. The AngleCam V2 performance assessment on the validation dataset resulted in an R^2^ = 0.62 and RMSE = 9.32° (n = 899, Fig. 3b).

**Figure 3:**
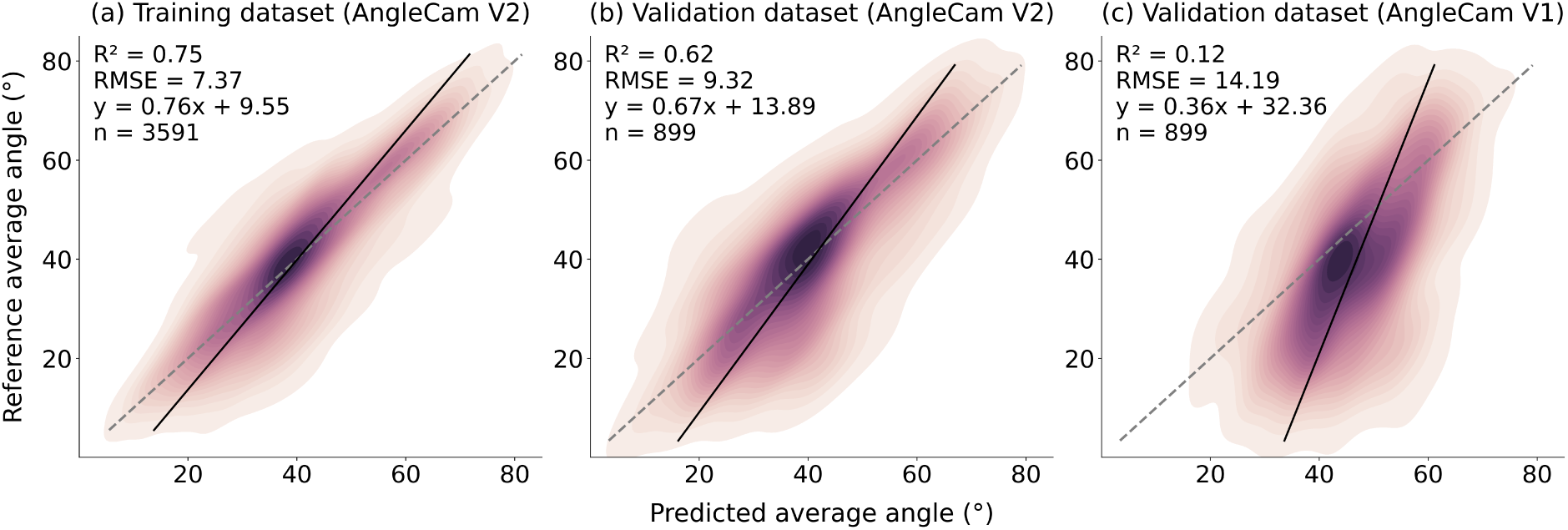
Comparison of AngleCam V1 and V2 performance across training and validation dataset. Kernel-density scatter plots show the joint distribution of predicted and reference average leaf angles (0–90°) for (a) the training set of AngleCam V2, (b) the independent validation set of AngleCam V2, and (c) the same validation set evaluated with AngleCam V1. Colors represent point density (dark = high density), the dashed grey line marks the 1:1 relationship, and the solid black line is the ordinary-least-squares regression fit (equation inset). Insets additionally report the coefficient of determination (R^2^), root-mean-square error (RMSE, degrees), and sample size (n).

Comparison with AngleCam V1 on the same validation dataset revealed clear performance improvements of AngleCam V2. AngleCam V1 achieved R^2^ = 0.12 with RMSE = 14.19° (Figure 3c). Its predictions exhibited greater dispersion and stronger systematic bias, tending to predict intermediate average leaf angles.

### 3.2 Phylogenetic error assessment

We analyzed prediction errors across 100 genera to test whether model performance was systematically structured by phylogeny. Mean absolute errors ranged from around 1° to 15°, with the majority of genera falling between 4° and 10° (Fig. 4). The smallest errors were observed in genera such as *Pilea* and *Nothofagus*, while the largest occurred in *Arbutus* and *Olea* (see Table A1 in the Appendix for all genus-level MAE values). When mapped onto the phylogenetic tree, prediction errors showed no significant clustering. Pagel’s *λ* was near zero (*λ* = 7.41 *×* 10*^−^*^5^), with no significant deviation from the null hypothesis of *λ* = 0 (no phylogenetic structure, *p* = 1.0), suggesting no phylogenetic structure in the residuals. Consistently, Moran’s *I* showed weak positive autocorrelation (*I* = 0.074, *p* = 0.23), supporting the finding that model residuals were not significantly structured by evolutionary relatedness.

**Figure 4:**
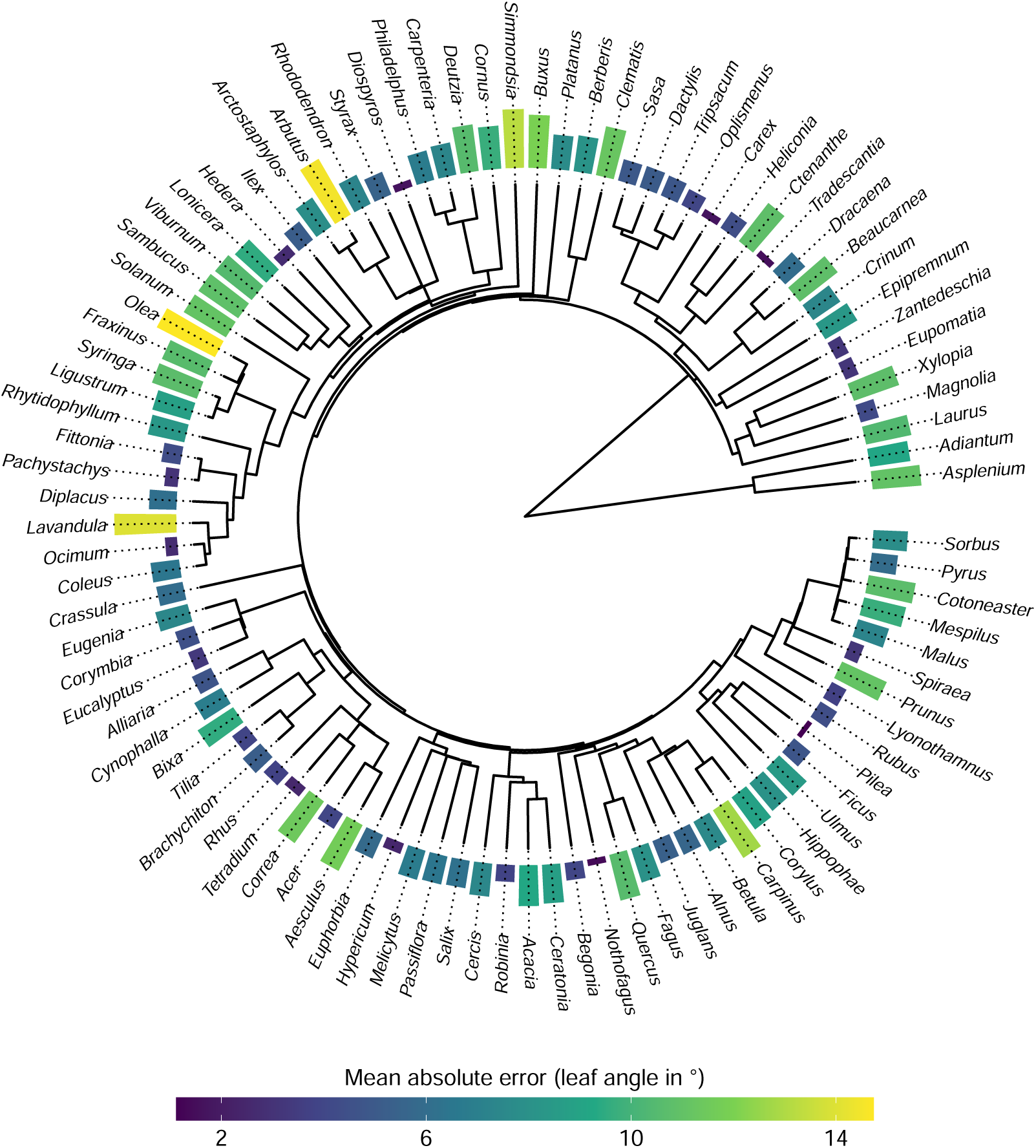
Phylogenetic distribution of AngleCam V2 prediction performance across 100 plant genera. Colored outer bars show residual per genus, computed for genera with at least three observations. Residuals show no significant clustering by clade. Pagel’s *λ* is near zero (*λ* = 7.41 *×* 10*^−^*^5^, *p* = 1.0), indicating no phylogenetic structure. Moran’s *I* shows weak autocorrelation (*I* = 0.074, *p* = 0.23). Together, these results indicate that there is no detectable phylogenetic autocorrelation in model errors.

### 3.3 Multitemporal model evaluation of AngleCam V2 with TLS-derived leaf inclination angle distributions

We validated AngleCam V2 predictions against independent TLS measurements using two indoor plant species (*Calathea ornata* and *Maranta leuconeura*) monitored continuously over nearly three days. Both species displayed pronounced diurnal leaf angle movements, which were captured by both the AngleCam and the TLS approach (Fig. 5).

**Figure 5:**
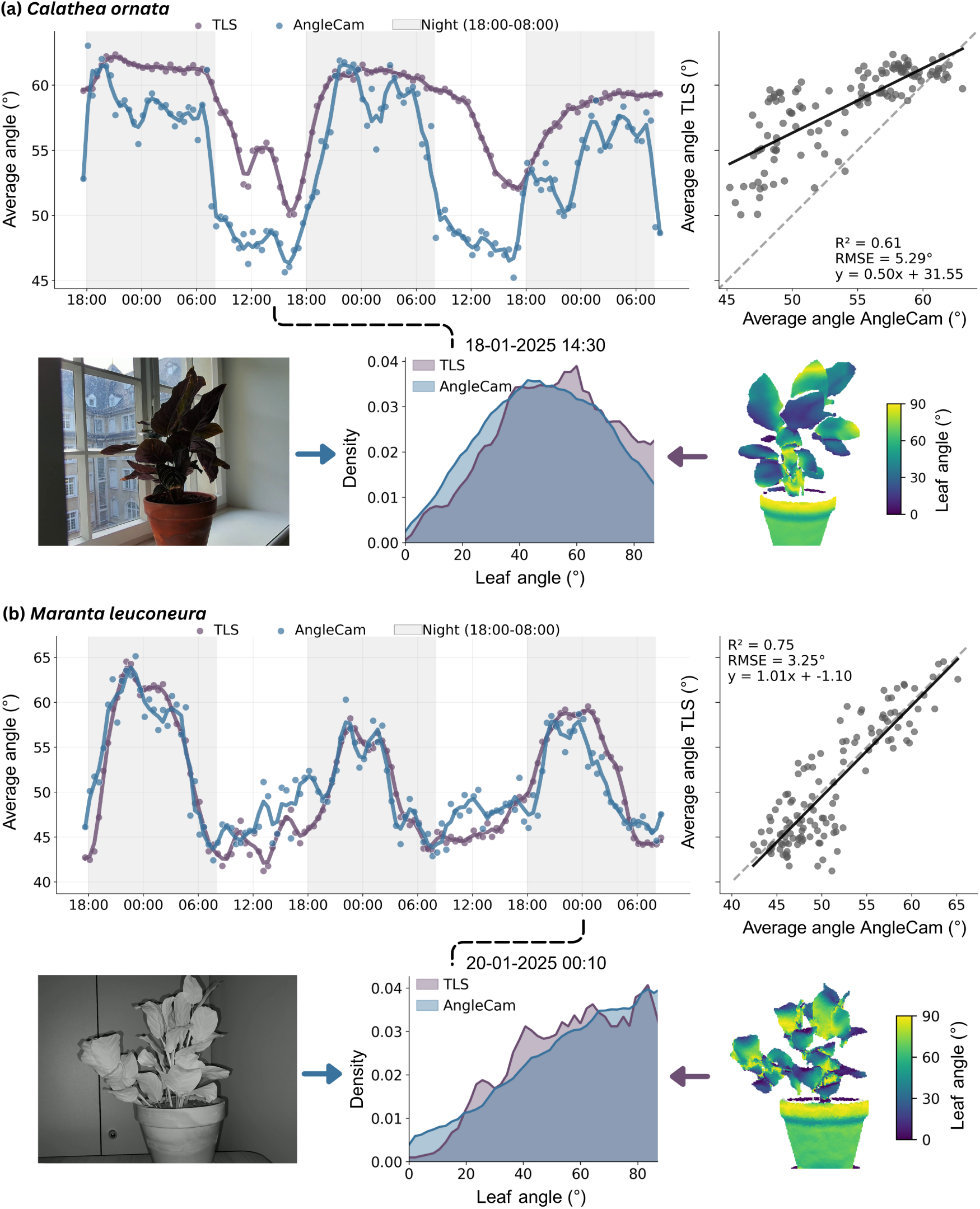
Multitemporal evaluation of AngleCam V2 predictions against TLS measurements for two plant species known for strong diurnal leaf movements. Time series show average leaf angles over nearly three days for (a) *Calathea ornata* (R^2^ = 0.61, RMSE = 5.29°) and (b) *Maranta leuconeura* (R^2^ = 0.75, RMSE = 3.25°), with both RGB (day) and near-infrared night-vision (night) imagery captured every 30 minutes. Scatter plots (right panels) show correspondence between methods across all time points. Distribution plots show example leaf inclination angle distributions at specific timestamps, with images illustrating plant appearance and 3D point cloud visualizations of the same setting. An interactive visualization can be assessed at Anonymous GitHub.

For *Calathea ornata*, AngleCam predictions followed the TLS trends, but there was a systematic discrepancy throughout the monitoring period, most pronounced at lower average leaf angles (Fig. 5a). The range of average leaf angles obtained by the two methods was very similar, ranging from approximately 45° to 63° for AngleCam and 50° to 63° for TLS. The average leaf angles derived from the two methods yielded a high correspondence with *R*^2^ = 0.61 and RMSE = 5.29°, with larger discrepancies for lower leaf angles and a closer fit at higher leaf angles.

The results for *Maranta leuconeura* showed stronger agreement (Fig. 5b). AngleCam predictions closely matched TLS-based estimates across the entire time series, with average angles ranging from approximately 42° to 65° for both methods. The correspondence was *R*^2^ = 0.75 and RMSE = 3.25°.

### 3.4 Multitemporal model evaluation of AngleCam V2 under water limitation

We assessed AngleCam’s sensitivity for tracking water limitation-induced changes in leaf angles by monitoring a non-watered individual of *Aglaonema commutatum* over an extended period of 14 days (Fig. 6a). The time series revealed a consistent increase in average leaf angle from approximately 43° to 48°, becoming especially evident after December 25. From this point onward, angles consistently exceeded baseline values and showed increased diurnal amplitude. The imagery (Fig. 6b) and corresponding leaf inclination angle distributions extracted at four time steps confirmed this trend, illustrating a gradual shift toward steeper angles (Fig. 6c). The overall shape of the distributions remained largely stable, with a slight tendency toward right skew, underscoring the robustness of AngleCam predictions under gradually changing physiological conditions.

**Figure 6:**
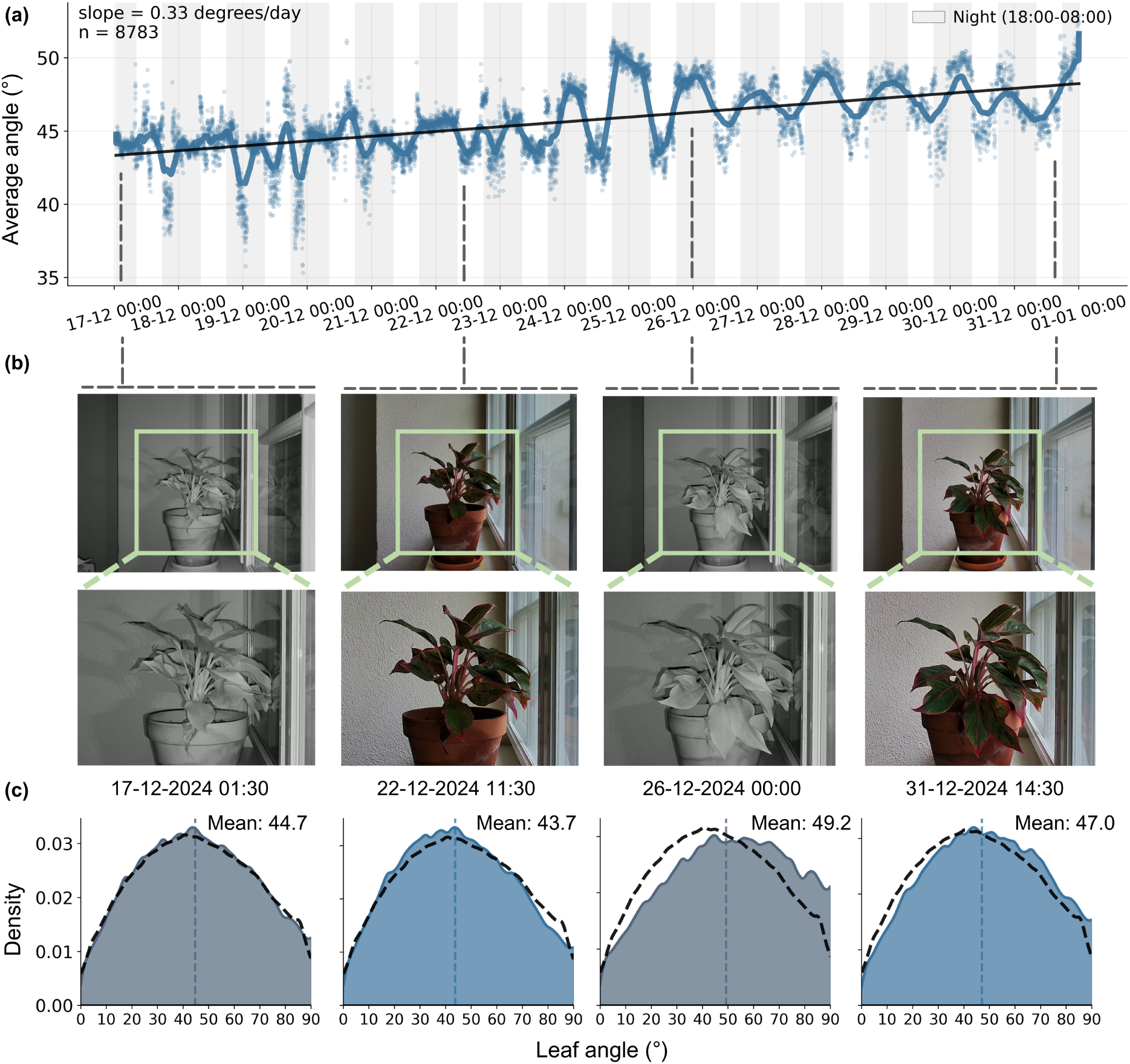
Tracking of leaf angle changes in *Aglaonema commutatum* induced by water limitation (14 days without irrigation). (a) Time series showing a gradual increase in average leaf angles from approximately 43° to 48° (slope = 0.33°/day, n = 8783 measurements), with pronounced diurnal oscillations and increased amplitude after December 25. The black line shows a linear trend with smoothed daily patterns overlaid. (b) Sequential NIR and RGB imagery at four time points showing visual progression of water limitation effects (c) Corresponding leaf inclination angle distributions at the same time points, demonstrating the gradual shift toward steeper angles (dashed line shows average distribution of all 8783 samples) while maintaining overall distribution shape. The results demonstrate the potential of AngleCam for revealing gradual physiological changes and its application for long-term drought monitoring.

## 4 Discussion

### 4.1 Model performance on training and validation data

Our evaluation revealed that AngleCam V2 achieved stronger predictive performance compared to its predecessor (AngleCam V1). When evaluated on their respective original validation datasets, AngleCam V1 showed higher performance metrics (validation: R^2^ = 0.84, RMSE = 6.13°; Kattenborn et al. (2022)) compared to AngleCam V2 (validation: R^2^ = 0.62, RMSE = 9.32°). However, when both models were evaluated on the new, more globally representative dataset, AngleCam V2 demonstrated substantially superior generalization, with an R^2^ of 0.62, compared to AngleCam V1’s R^2^ of 0.12. This increased performance can be mainly attributed to the fact that AngleCam V2 was trained on an expanded dataset with considerably greater species diversity and scene heterogeneity than the first version. This expansion deliberately introduced more variation in leaf morphologies, growth forms, environmental conditions, cameras, and data acquisition settings, enhancing the model’s ability to generalize across diverse ecological contexts and data acquisition scenarios.

The expanded dataset inevitably introduced additional label noise as well as variability in image features unrelated to actual leaf angle differences, both of which can bias a model (Belkin et al., 2019; Geman et al., 1992). While large training sets can help mitigate such effects (Rolnick et al., 2017), complex deep learning models may still memorize noise or spurious patterns instead of extracting meaningful signals. Given our dataset size and the deep transformer model (DINOv2), this risk would manifest as low bias but high variance if regularization were insufficient. To address this, we implemented stronger regularization strategies compared to AngleCam V1. A dropout rate of 0.4 in the regression head randomly deactivated 40% of units during each forward pass, facilitating the model to rely on robust features rather than memorizing training-specific patterns. Weight decay provided additional regularization by penalizing large parameter values, encouraging smoother angle approximations (Loshchilov & Hutter, 2017; Zhang et al., 2018). We also transitioned from mean squared error (MSE) to Huber loss, which treats outliers more robustly by being quadratic for small errors and linear for large errors (Sun et al., 2020). Together, these techniques facilitated the model to extract meaningful signals from the expanded dataset while ensuring the generalization of the model over the more variable scene conditions and taxa.

The validation results revealed a tendency where low leaf angles were overestimated and high angles were underestimated. We assume that this bias, also known as regression to the mean bias (Barnett et al., 2005), emerges from the training data distribution, which contains abundant samples between 35°-50° but relatively few extreme cases below 20° or above 60°. Several approaches could address this imbalance, such as loss weighting to give more importance to rare, extreme angles during training, or a weighted training sampler that increases the selection frequency of underrepresented angle ranges. However, such interventions require careful implementation to avoid introducing artificial biases or unstable prediction behavior (Barnett et al., 2005). AngleCam development remains an ongoing process, and future versions will focus on expanding training diversity in extreme angle scenarios.

### 4.2 Phylogenetic error assessment

The residual analysis revealed no statistically significant phylogenetic signal in prediction errors (Fig. 4), indicating that AngleCam V2 learned general principles of leaf angle estimation that transfer across taxonomic and evolutionary lineages. The absence of phylogenetic autocorrelation suggests the model captures fundamental geometric relationships rather than taxon-specific visual features. Pagel’s *λ* was near zero (*λ* = 7.41 *×* 10*^−^*^5^, *p* = 1.0), indicating no phylogenetic structure in prediction errors. Similarly, Moran’s *I* showed only weak, non-significant autocorrelation (*I* = 0.074, *p* = 0.23), confirming the lack of systematic phylogenetic bias.

The substantial representation of *Tilia*, *Acer*, *Quercus*, and *Fagus* in our training dataset (see also Fig. 1, Tab. A1) results from continuous time-series monitoring, providing extensive environmental and phenological coverage across these genera. Notably, the phylogenetic analysis demonstrates that prediction errors are not systematically lower for such well-represented genera compared to those with fewer training samples. Although more balanced taxonomic representation would be ideal for future datasets, the temporal richness within these dominant genera added training diversity across environmental conditions without compromising model generalization.

This supports applying AngleCam to new species, given that their broader taxonomic groups are reasonably represented in the training data. Still, generalization has its limits: performance may degrade for clades that are sparsely or not represented at all, and clade-specific biases may emerge in such cases. Overall, the lack of systematic phylogenetic bias suggests that within-genus variation in leaf morphology and growth dynamics is sufficiently captured by our training set, allowing for reasonable predictions for closely related species.

### 4.3 Evaluating AngleCam on representing leaf angle diurnal dynamics

Our controlled testing experiments with TLS demonstrated AngleCam V2’s ability to track temporal leaf angle dynamics across day-night cycles. The successful tracking of continuous diurnal patterns represents a significant advancement. As demonstrated for *Maranta leuconeura* and *Calathea ornata*, the model captured natural circadian leaf movements despite never being explicitly trained on labeled NIR imagery. This capability emerged from our dual-modality training strategy, combining grayscale conversion with pseudo-NIR generation. As shown by the TLS comparison across the day and night cycles, this strategy proved highly effective. It avoided the need for additional NIR labeling, reduced annotation effort, and at the same time harnessed the full potential of the extensive RGB dataset already available. However, this dual-modality approach may slightly reduce RGB-specific performance compared to a purely RGB-trained model. Future versions incorporating independently labeled NIR images could enhance both RGB and night-vision capabilities.

The systematic underestimation observed for *Calathea ornata* relative to TLS measurements may be partly attributed to sampling biases inherent to the laser scanning methodology. While horizontally oriented leaf surfaces have a higher chance of being occluded, more vertically oriented leaves intercept more laser pulses. This generates higher point densities that can inflate average angle estimates toward steeper values (Jiang et al., 2021). While our voxel downsampling to 1 cm resolution reduced this effect, some bias likely remained. This sampling artifact was less pronounced for *Maranta leuconeura* due to its smaller leaf dimensions and overall more spherically oriented leaf inclination angle distributions (Fig. 5).

Species-specific performance variations may also indicate current model generalization limits. The superior performance for *Maranta leuconeura* (R^2^ = 0.75, RMSE = 3.25°) compared to *Calathea ornata* (R^2^ = 0.61, RMSE = 5.29°) may reflect differences in training data representation. Both species had RGB samples from the experimental setup, acquired several months prior, included in the training data. However, *Maranta leuconeura* benefited from the presence of morphologically similar species, particularly *Ctenanthe burle-marxii* (n = 7), which share comparable leaf architecture and growth patterns. In contrast, *Calathea ornata* lacked morphologically similar representatives beyond the experimental samples themselves. This indicates that model performance can depend on exposure to similar species during training. When deploying AngleCam for experimental monitoring, particularly for species that are not adequately represented by morphologically similar taxa in the training dataset, we recommend acquiring at least 15-20 labeled images from the target scene for model calibration. We provide an open-source labeling tool, pre-trained model weights, and a retraining framework (Anonymous GitHub), enabling a flexible adaptation of AngleCam to particular experimental conditions.

### 4.4 Application of AngleCam in future research

AngleCam V2 addresses limitations in low-cost and scalable leaf angle monitoring by enabling continuous day-night observations, opening up new possibilities for ecosystem research. The method’s compatibility with standard time-lapse cameras, low hardware costs relative to laser scanning systems, and minimal field maintenance requirements make it a valuable asset for long-term monitoring networks (e.g., flux-towers, biodiversity experiments, PhenoCam). Integration with eddy covariance measurements could provide insights into how diurnal and seasonal leaf movements are coupled with carbon fluxes and ecosystem-atmosphere interactions (Baldocchi, 2020). Given that variable vertical leaf angle profiles impact light interception and canopy energy balance, continuous monitoring could enhance our understanding of ecosystem responses to environmental changes (Yang et al., 2023).

Despite recent advances, knowledge about factors driving leaf angle dynamics remains limited. While diurnal patterns and vertical LIAD gradients are documented (Ehleringer & Forseth, 1980; Kao & Forseth, 1992; Liu et al., 2019; Pisek et al., 2022), the relative importance of light availability versus water stress in controlling these dynamics is poorly understood (Kattenborn et al., 2022; Yang et al., 2023). AngleCam’s high temporal resolution and multi-height deployment capability canhelp to address this knowledge gap. Over the course of the 14-day experiment in this study, progressive soil drainage (with the plant remaining unwatered for a total of 19 days) revealed that the leaf inclination angle distributions of *Aglaonema commutatum* responded rapidly to water limitation. AngleCam’s sensitivity to changes in plant condition enabled quantification of these dynamics, demonstrating its potential for tracking plant responses to climate extremes, such as atmospheric or soil drought. Interestingly, we observed two trends: a steady increase in average leaf angles and a progressive amplification of diurnal variation. These findings align with a previous study on five temperate tree species, which showed that leaf angle dynamics respond both in absolute values and in their temporal variability (Kattenborn et al., 2022, 2024). In the future, simultaneous monitoring of plant physiological parameters (e.g., sap flow, stem water potential), environmental conditions (e.g., temperature, light intensity), and ecosystem fluxes (e.g., carbon and water exchange) alongside leaf angle dynamics could help uncover the mechanistic drivers of these dynamics and support the use of leaf angle as an indicator of plant physiological status. However, given that species respond differently to environmental stresses, such applications would likely require species-specific training and calibration to establish reliable relationships between leaf angle changes and physiological conditions. Past and current deployments of AngleCam reflect its potential for ecological and plant physiological research. Installations at the Leipzig Canopy Crane (Richter et al., 2021), the MyDiv experiment (Ferlian et al., 2018), and the ArboFun experiment (Kretz et al., 2025), all part of the German Centre for Integrative Biodiversity Research (iDiv), explore how structural, taxonomic, and functional diversity shape leaf angle dynamics for temperate tree species. In addition, AngleCam installations at the ECOSENSE site in Ettenheim (Werner et al., 2024) and at the Hartheim Research Station in Germany (Integrated Carbon Observation System (ICOS) flux-tower site) cover multiple individuals of temperate broad-leaved tree species. All of the above-mentioned installations capture leaf movements alongside comprehensive physiological and environmental measurements. Expanding such integrated monitoring to other research infrastructures and biomes could significantly enhance our understanding of leaf angle dynamics. Existing PhenoCam and eddy covariance flux tower networks (Brown et al., 2016; Richardson et al., 2018) offer ideal platforms for scaling up AngleCam installations and linking leaf angle variability to plant and ecosystem function at broader spatial and temporal scales.

Beyond ecological and physiological monitoring at local scales, understanding leaf angle dynamics is also critical for interpreting vegetation signals in satellite-based Earth observation. Leaf angles affect the scattering and absorption of light within plant canopies and thereby influence radiative transfer processes (Hase et al., 2022; Kattenborn et al., 2019). Temporal changes in vertical leaf angles can significantly alter reflectance signals and systematically confound widely used vegetation indices, such as kNDVI (Hase et al., 2022; Kattenborn et al., 2024) or fluorescence signals (Jablonski et al., 2025). These changes are often not random noise but reflect structured responses to environmental drivers like temperature or radiation (Kattenborn et al., 2022). Thus, unresolved leaf angle dynamics may obscure or mimic other physiological variability, such as vegetation productivity changes when observed from space. At the same time, this sensitivity to environmental cues means that remotely sensed variations in canopy reflectance may contain embedded signals of leaf angle movements. AngleCam can thus contribute to disentangling these effects by providing temporally resolved parameters in radiative transfer modeling or by acting as a covariate to interpret better satellite observations of vegetation stress (Hase et al., 2022; Jablonski et al., 2025; Kattenborn et al., 2024).

The model’s demonstrated generalization across different scene conditions, growth forms, and taxonomic and phylogenetic lineages can unlock large-scale applications in concert with citizen science platforms. Particularly, citizen science apps for plant species recognition, such as iNaturalist or Pl@ntNet, acquire millions of geolocation plant images each year, many of which are acquired with sufficient distance and camera perspectives compatible with AngleCam (Garcin et al., 2021; Mason et al., 2025; Soltani et al., 2024). Previous studies have already highlighted the value of citizen science data for uncovering plant functional traits or phenology across spatiotemporal scales (Mora et al., 2024; Schiller et al., 2021; Wolf et al., 2022). Accordingly, citizen science data may also represent an unprecedented data treasure to uncover global patterns in leaf inclination angle distributions at the species and community level, biogeographic variations in leaf positioning strategies, or seasonal adaptation patterns across biomes. This approach would substantially expand our understanding of how leaf angles are organized within and across species worldwide, potentially uncovering previously unknown global patterns in plant structural adaptation (Li & Fang, 2024).

## 5 Conclusions

We introduced AngleCam V2, an enhanced deep learning model for estimating leaf inclination angle distributions from RGB and night-vision NIR imagery. By expanding training data across species, canopy structures, and lighting conditions, as well as integrating synthetic near-infrared augmentation, the model shows improved generalization and temporal robustness, including nighttime compatibility.

Testing against terrestrial laser scanning (TLS) confirmed that AngleCam reliably tracks LIADs across day and night imagery. As demonstrated with a water limitation experiment, AngleCam could reveal multi-day trends in LIADs variability. Furthermore, phylogenetic analysis revealed no systematic prediction bias across taxa, supporting its generalizability across a wide range of plant species.

A central feature of AngleCam is its broad applicability: it can be used with standard imagery from smartphones and time-lapse cameras to PhenoCam networks, provided cameras are oriented horizontally. The method requires no specialized hardware, making it easy to deploy in field, lab, or citizen science settings. This enables both high-frequency monitoring of leaf angle dynamics over time and flexible snapshot-based assessments using handheld devices. AngleCam V2 thus provides a scalable, accessible, and open-source tool for monitoring leaf angle dynamics across time, taxa, and environments.

## Acknowledgements

LK and TK received further funding from the German Research Foundation (DFG) under the projects PANOPS (project number 504978936) and ECOSENSE (project number SFB 1537/1). TK received funding from the Flexible Funds Program for junior scientists of the University of Leipzig (project number 232201582). TK and CW gratefully acknowledge the support of iDiv funded by the German Research Foundation (DFG–FZT 118, 202548816). JP was supported by the Estonian Research Council Grant PRG 1405 and the Estonian Ministry of Education and Research, Centre of Excellence for Sustainable Land Use (TK232).

## Author Contributions

LK and TK conceived the ideas, designed the methodology, and led the analysis. TK, JP, RR, JF, LK, and DL collected the data. LK and TK led the writing of the manuscript. All authors contributed critically to the drafts and gave final approval for publication.

## Data Availability Statement

The code is available here (https://anonymous.4open.science/r/AngleCamV2-2B38). The data is available at (https://doi.org/10.5281/zenodo.17086253). The pretrained model is available at (https://doi.org/10.5281/zenodo.17101166).

## A Appendix

### A.1 Genus representation across datasets

**Table A1:**
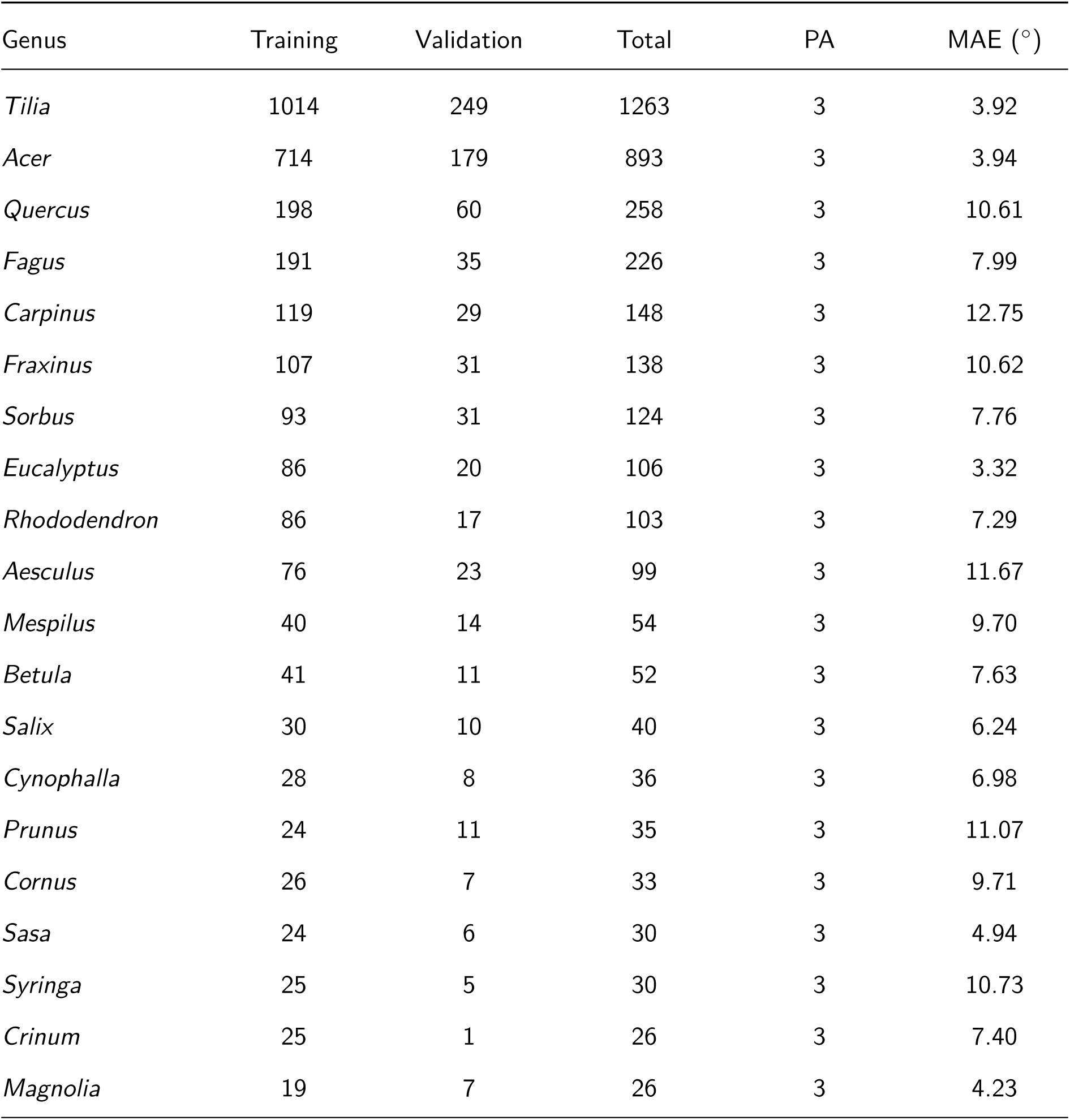
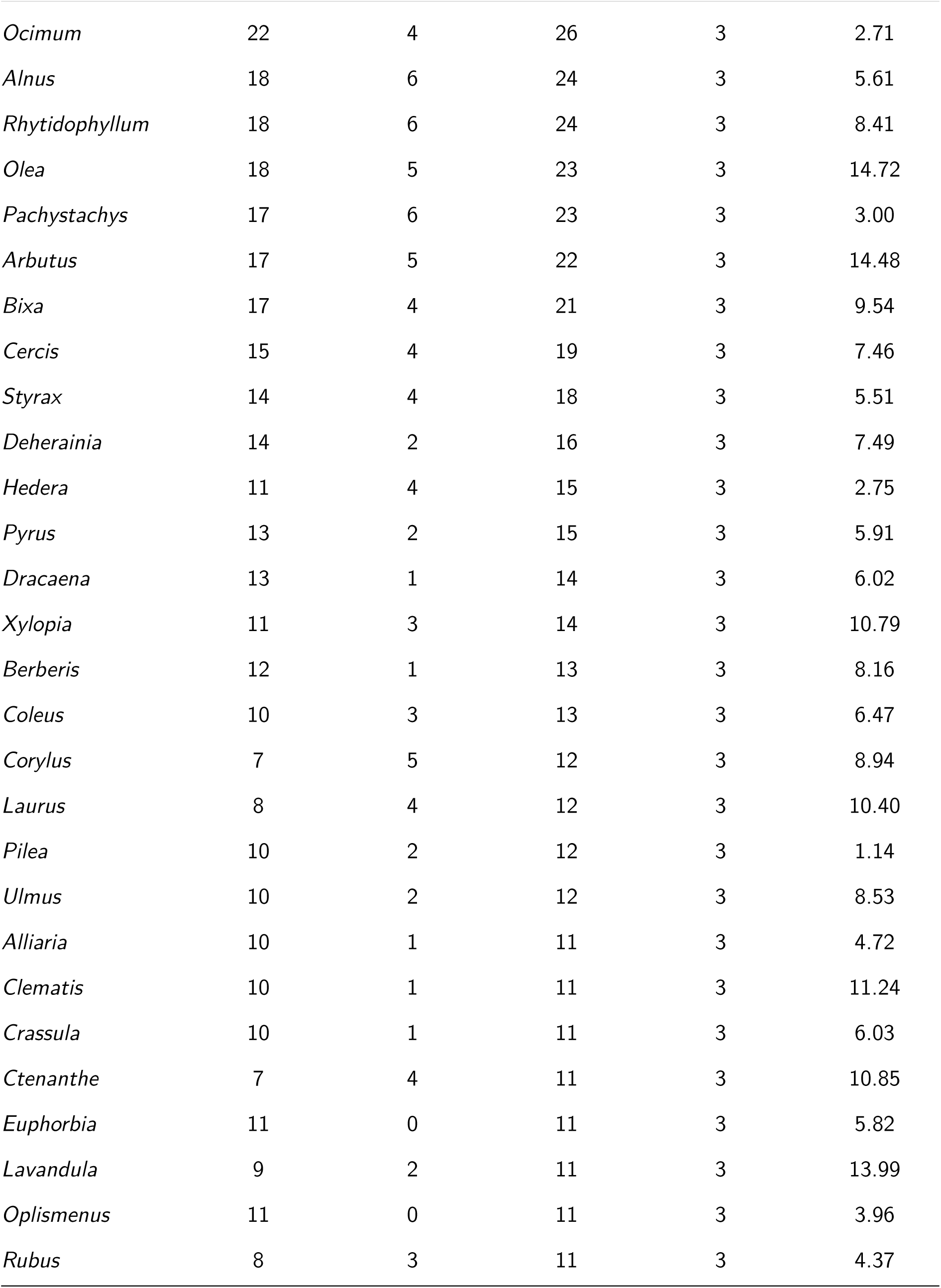
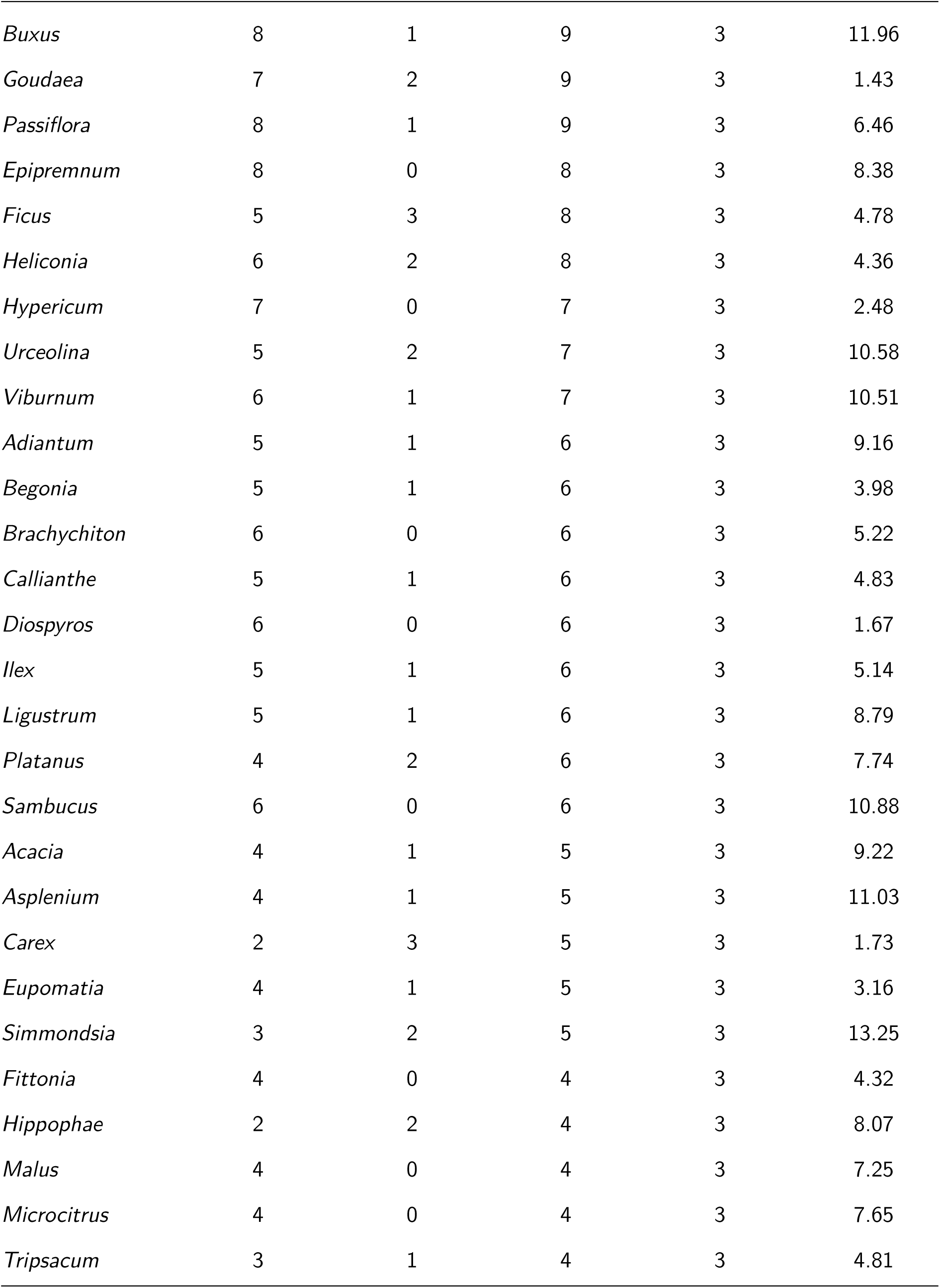
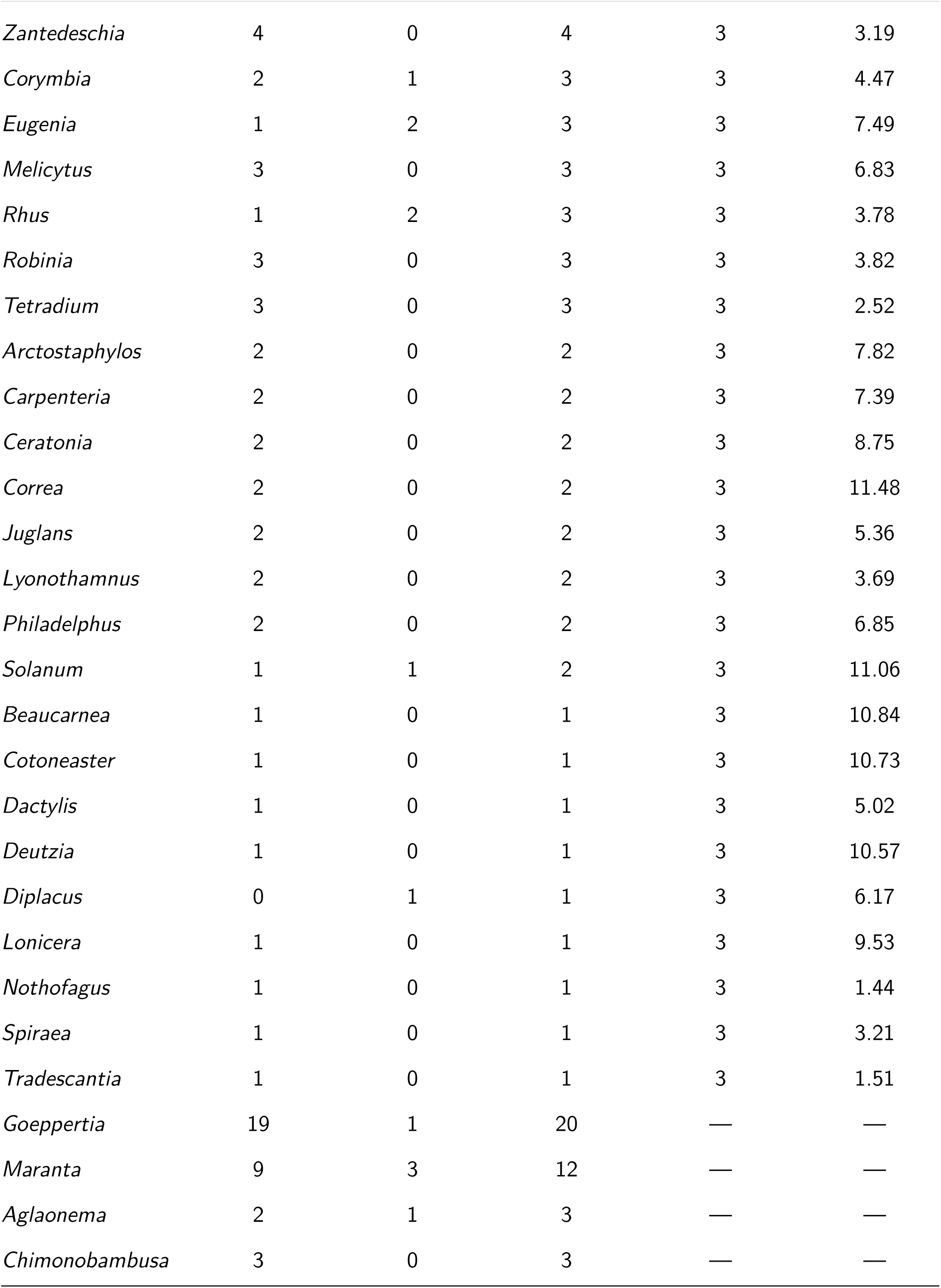
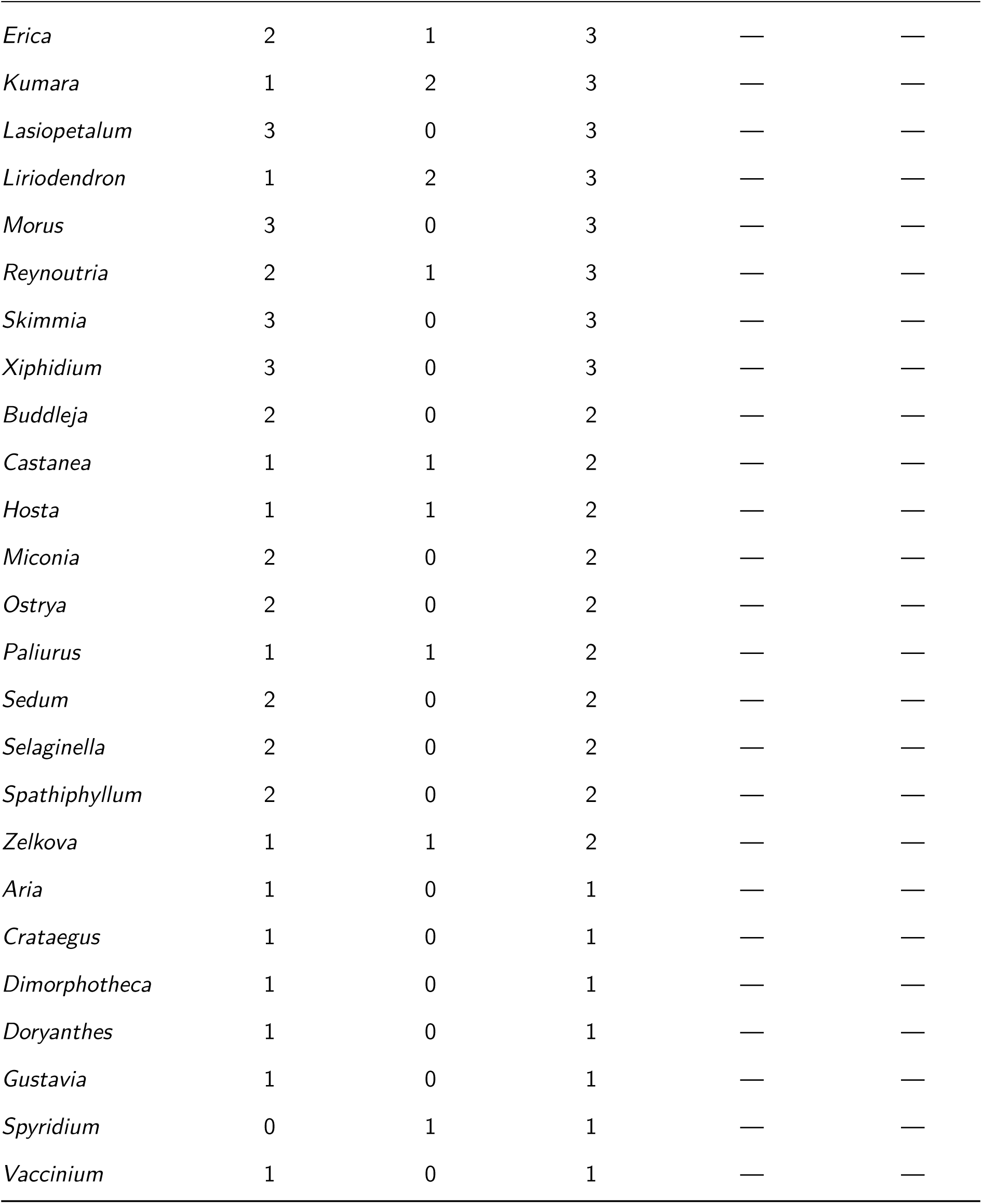
Genus-level representation across training, validation, and phylogenetic analysis (PA) datasets. Training and validation counts show the number of images per genus used for model development. Phylogenetic analysis includes three samples per genus where available, with mean absolute error (MAE) values representing the mean absolute deviation between AngleCam V2 predictions and labeled leaf inclination angle distribution references in degrees. Empty MAE values (—) indicate genera not included in the phylogenetic analysis. Genera are ordered by phylogenetic data availability and total sample count.

